# HKDC1 contributes to aberrant lysosome-mitochondria contact in Niemann-Pick disease type C

**DOI:** 10.64898/2026.06.01.729255

**Authors:** Raffaele Pastore, Jack Stanley, Szilvia Kiraly, Christelle Dubois, Jlenia Monfregola, Antonio De Luca, Germano Guerra, Mark Skehel, Andrea Ballabio, Frances M. Platt, Emily R. Eden

## Abstract

Niemann-Pick disease type C (NPC) is a neurovisceral lysosomal storage disorder comprising two clinically indistinguishable but genetically distinct subtypes caused by mutations in *NPC1*, or *NPC2*. The specific impact of each deficiency on cellular homeostasis remains poorly defined due to the phenotypic heterogeneity of patient-derived models and a lack of isogenic platforms for comparative study. Here we established isogenic ARPE19 models of *NPC1* and *NPC2* deficiency that faithfully recapitulate hallmark pathologies, including homogeneous lysosomal expansion and lipid sequestration. Direct comparison of these isogenic lines revealed a fundamental divergence in organelle crosstalk: while both genotypes exhibit comparable lipid accumulation, expanded mitochondria-lysosome contact sites (MLCs) are observed exclusively in *NPC1^−/−^* cells. Using StARD3-targeted proximity labelling and quantitative proteomics, we identified the mitochondrial protein HKDC1 as an MLC regulator. We demonstrate that HKDC1 is markedly upregulated in *NPC1^−/−^* cells and that its overexpression drives MLC expansion in wild-type cells. Thus our study uncovers a homeostatic role for HKDC1-mediated organelle remodelling and demonstrates the power of isogenic modelling for identifying novel regulators of organelle architecture and potential therapeutic targets.

## Introduction

Niemann-Pick disease type C (NPC) is a rare autosomal recessive lysosomal storage disorder caused by mutations in either *NPC1* or *NPC2*. These two genes encode proteins that cooperate in the trafficking of cholesterol and sphingolipids from late endosomes/lysosomes (Kwon et al., 2009) (Qian et al., 2020). Loss of function in either protein leads to abnormal accumulation of unesterified cholesterol, sphingosine, and sphinganine, accompanied by secondary defects in glycosphingolipid trafficking, lysosomal function, cellular signaling and the dynamics of organelle membrane contact sites (Lloyd-Evans et al., 2008) (Höglinger et al., 2019); (Pastore et al., 2025). NPC patients present with a spectrum of neurological, hepatic, and pulmonary manifestations, with *NPC1* mutations accounting for the majority of cases and *NPC2* mutations being far less common but clinically indistinguishable (Vanier, 2015). There has been considerable progress in identifying pathogenic variants and three small molecule disease modifying therapies are now approved. Until 2024, the glucosylceramide synthase inhibitor miglustat was the only approved drug and has significantly extended lifespan (Patterson et al., 2020). The therapeutic landscape expanded in 2024, with the FDA approvals of the new drugs *N*-acetyl-L-leucine (Aqneursa) and arimoclomol (Miplyffa), the latter indicated for use in combination with miglustat (Klein et al., 2025). While these approvals mark major and important progress, the long-term therapeutic benefit and the ability to achieve disease modification across all NPC patients including NPC2 remains unclear, highlighting the need for improved mechanistic models to elucidate disease pathophysiology and accelerate drug discovery.

One of the main limitations in NPC research has been the lack of robust cellular models that permit direct comparisons between NPC1 and NPC2 deficiency. Patient-derived fibroblasts and induced pluripotent stem cells (iPSCs) display inconsistent phenotypes and high variability, while iPSCs are technically demanding, time consuming, biochemically variable and costly to maintain (Vanier et al., 1996) (Tängemo et al., 2011), (Prabhu et al., 2021). Knockdown models, such as shRNA-mediated systems, often produce incomplete gene suppression and heterogeneous results (Rodríguez-Pascau et al., 2012). Pharmacological inhibition of NPC1 with U18666A which interacts with the sterol sensing domain (SSD) of NPC1 is a useful and widely used tool (Liscum & Faust, 1989), but has also been implicated in inhibition of other lipid transport proteins including GRAMD proteins (Xiao et al., 2021) (Lu et al., 2015). Furthermore, existing models are usually restricted to either NPC1 or NPC2, preventing direct comparisons of the two disease subtypes in the same genetic background (Pallottini & Pfrieger, 2020). This is a critical gap, as it remains unclear whether NPC1 and NPC2 loss of function results in the same or subtly distinct cellular phenotypes.

To address these limitations, we developed novel isogenic human cellular models of NPC by generating *NPC1*^⁻/⁻^ and *NPC2*^⁻/⁻^ knockouts in the ARPE19 human retinal pigment epithelial cell line using CRISPR/Cas9-mediated genome editing (Jinek et al., 2012). These lines share an identical genetic background, thereby enabling for the first time, direct comparative studies of NPC1 and NPC2 deficiencies. Here, we report that while both models reproduce hallmark NPC cellular phenotypes, they reveal fundamental divergence in organelle crosstalk, with expanded mitochondria-lysosome contact sites (MLCs) observed exclusively in *NPC1^−/−^* cells. We have previously shown that the late endosome/lysosome (LE/Lys) sterol transfer protein StARD3 (StAR-related lipid transfer domain-containing protein 3) drives the expansion of mitochondria-lysosome contacts (MLCs) upon deficiency of the NPC1 protein (Höglinger et al., 2019). StARD3 acts as a multi-valent structural tether capable of establishing distinct membrane contact sites (MCSs). Whereas in models of NPC pathology, upregulated StARD3 localizes to expanded MLCs (Höglinger et al., 2019), under normal physiological conditions, StARD3 instead tethers ER-endosome MCSs via interaction of its FFAT (two phenylalanines in an acidic tract) motif with ER-resident VAP and MOSPD2 proteins (Alpy et al., 2013; Eichler et al., 2026), facilitating direct, non-vesicular cholesterol transport through its START domain (Wilhelm et al., 2017). Leveraging StARD3’s inherent tethering activity, we used StARD3-targeted proximity labeling and quantitative proteomics to map the molecular landscape of expanded MLCs in NPC. This approach identified the mitochondrial protein hexokinase domain containing protein-1 (HKDC1) as a novel MLC regulator. We demonstrate that HKDC1 is markedly upregulated in *NPC1^−/−^* cells and that its overexpression is sufficient to drive MLC expansion in wild-type cells, establishing a direct mitochondrial contribution to NPC1 pathology. These findings highlight the potential of NPC ARPE19 isogenic cell models as robust tools for dissecting NPC pathophysiology and accelerating drug discovery.

## Results

### Loss of NPC1 or NPC2 causes lysosomal enlargement and lipid trafficking defects

To investigate the roles of NPC1 and NPC2 in a consistent genetic background, we generated isogenic ARPE19 knockout lines using CRISPR/Cas9-mediated genome editing targeting exon 1 of *NPC1* or exon 4 of *NPC2* (Kim et al., 2017; Mali et al., 2013). Gene disruption was confirmed by RT-PCR, Western blotting, and immunofluorescence, demonstrating complete loss of the respective proteins (Fig. 1A-D, Supp. Fig. S1A-D). Both *NPC1^⁻/⁻^* and *NPC2^⁻/⁻^* ARPE19 cells exhibited pronounced lysosomal abnormalities that recapitulate key cellular features of Niemann-Pick disease type C (NPC) (Sun et al., 2001) (te Vruchte et al., 2014) (Las Heras et al., 2023) (Best et al., 2025). LAMP1 staining revealed perinuclear clustering of enlarged LE/Lys, in contrast to the dispersed distribution observed in wild-type (WT) ARPE19 cells (Fig. 2A, Supp. Fig. S3A). Similarly, LysoTracker fluorescence was elevated approximately three-fold in both knockout lines (Fig. 2B) and western blotting showed increased LAMP1 protein levels (Supp. Fig. S2A, B). Electron microscopy confirmed expansion of LE/Lys containing electron-dense multilamellar bodies, characteristic of lipid storage disorders (Fig. 2C, D) (Platt et al., 2018) (Katona et al., 2014) (Calcagni et al., 2023). Similar abnormalities were observed in patient-derived fibroblasts (Fig. S2C), although lysosomal size was highly variable both within and between patient lines, whereas ARPE19 knockout cells displayed a uniform NPC-like phenotype that enabled quantitative analysis (Fig. 2D).

**Figure 1:**
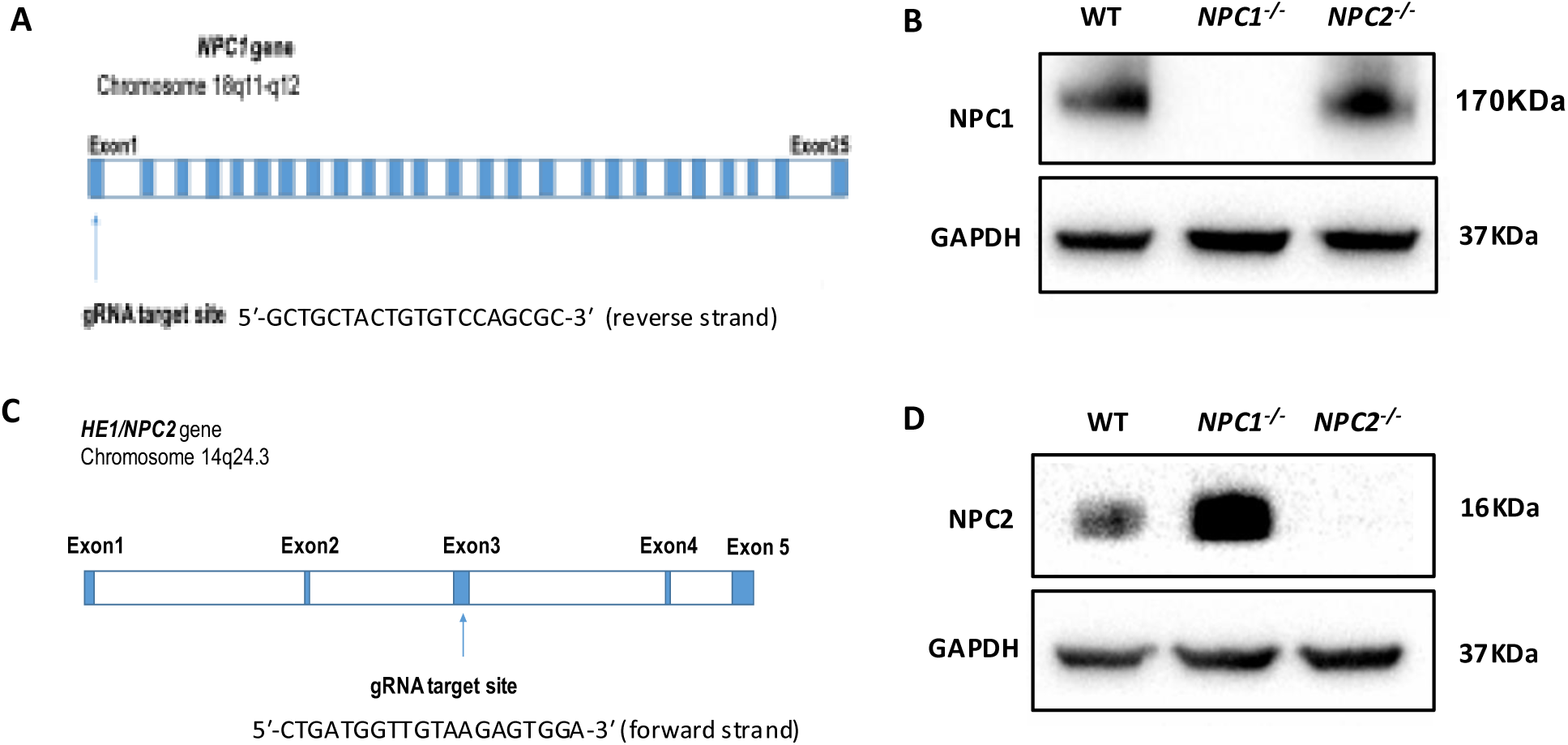
Generation and validation of ARPE-19 *NPC1^−/−^* and ARPE-19 *NPC2^−/−^* cell lines. (A, C) Schematic overview of CRISPR/Cas9-mediated targeting of the *NPC1* and *NPC2* genes in ARPE-19 cells. *NPC1* gene sgRNA target site: 5′-GCTGCTACTGTGTCCAGCGC-3′ (reverse strand). *NPC2* gene sgRNA target site: 5′-CTGATGGTTGTAAGAGTGGA-3′ (forward strand). (B, D) Western blot analysis of NPC1 and NPC2 protein levels in wild-type ARPE-19, ARPE-19 *NPC1^−/−^* and ARPE-19 *NPC2^−/−^*. GAPDH served as a loading control.

**Figure 2:**
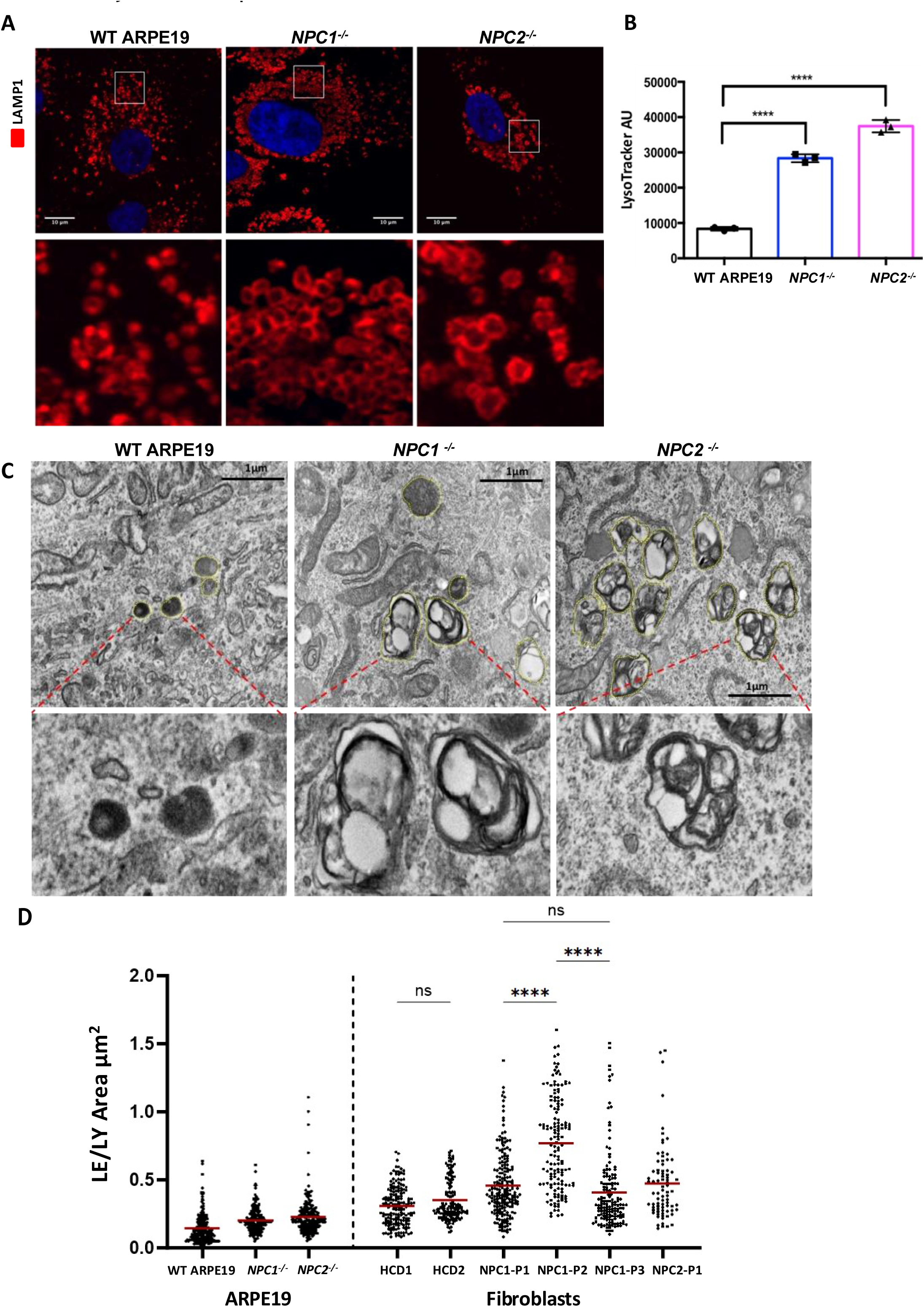
Lysosome expansion in ARPE-19 *NPC1^−/−^* and ARPE-19 *NPC2^−/−^* cells. (A) Confocal microscopy images of wild-type (WT), *NPC1*⁻/⁻, and *NPC2*⁻/⁻ ARPE-19 cells immunostained for LAMP1 (red) and counterstained with DAPI (blue). Scale bar: 10 µm. (B) Quantification of LysoTracker Green fluorescence intensity in the indicated cell lines measured by flow cytometry (200 nM LysoTracker, 10 min). A total of 10,000 events were recorded per sample. AU, arbitrary units. Data represent mean ± SD from n = 3 biological replicates. Statistical significance was assessed by one-way ANOVA followed by Dunnett’s multiple comparisons test; ****p < 0.0001. (C) Transmission electron microscopy (TEM) images of WT, *NPC1*⁻/⁻, and *NPC2*⁻/⁻ ARPE-19 cells. LE/LY are outlined with yellow dashed lines. Insets show enlarged lysosomes containing lipid vesicles and electron-dense multilamellar bodies. Scale bar: 1 µm. (D) Quantification of late endosomes/lysosomes (LE/LY) area (µm²) from TEM images of ARPE-19 WT, *NPC1^−/−^*, *NPC2^−/−^*, and human skin primary fibroblasts (Fig. S3) derived from healthy control donors (HCD) or NPC patients. ns=not significant. *p*-values were determined using Kruskal-Wallis followed by Dunn’s multiple comparisons test, \*\*\*\**p* < 0.0001.

These morphological changes correlated with hallmark lipid storage phenotypes. Biochemical analyses revealed an approximately three-fold increase in sphingosine, sphinganine, and total cholesterol in both knockout lines relative to WT ARPE19 cells (Fig. 3A-C). Confocal imaging showed that unesterified cholesterol accumulated specifically within LAMP1-positive compartments, as confirmed by filipin-LAMP1 co-staining (Fig. 3D, E; Supp. Fig. S3A). Primary patient fibroblasts exhibited substantial variability in filipin staining (Fig. 3F, G), whereas ARPE19 knockout cells provided a uniform and reproducible readout. Lipid accumulation was accompanied by impaired glycosphingolipid trafficking, a characteristic of sphingolipid disorders (Chen et al., 1999) (Sun et al., 2001), (Vruchte et al., 2004). In both *NPC1^⁻/⁻^* and *NPC2^⁻/⁻^* cells, BODIPY-labeled lactosylceramide (BODIPY-LacCer) appeared as dispersed cytoplasmic puncta, in contrast to the perinuclear Golgi localization observed in WT cells (Supp. Fig. S3B).

**Figure 3:**
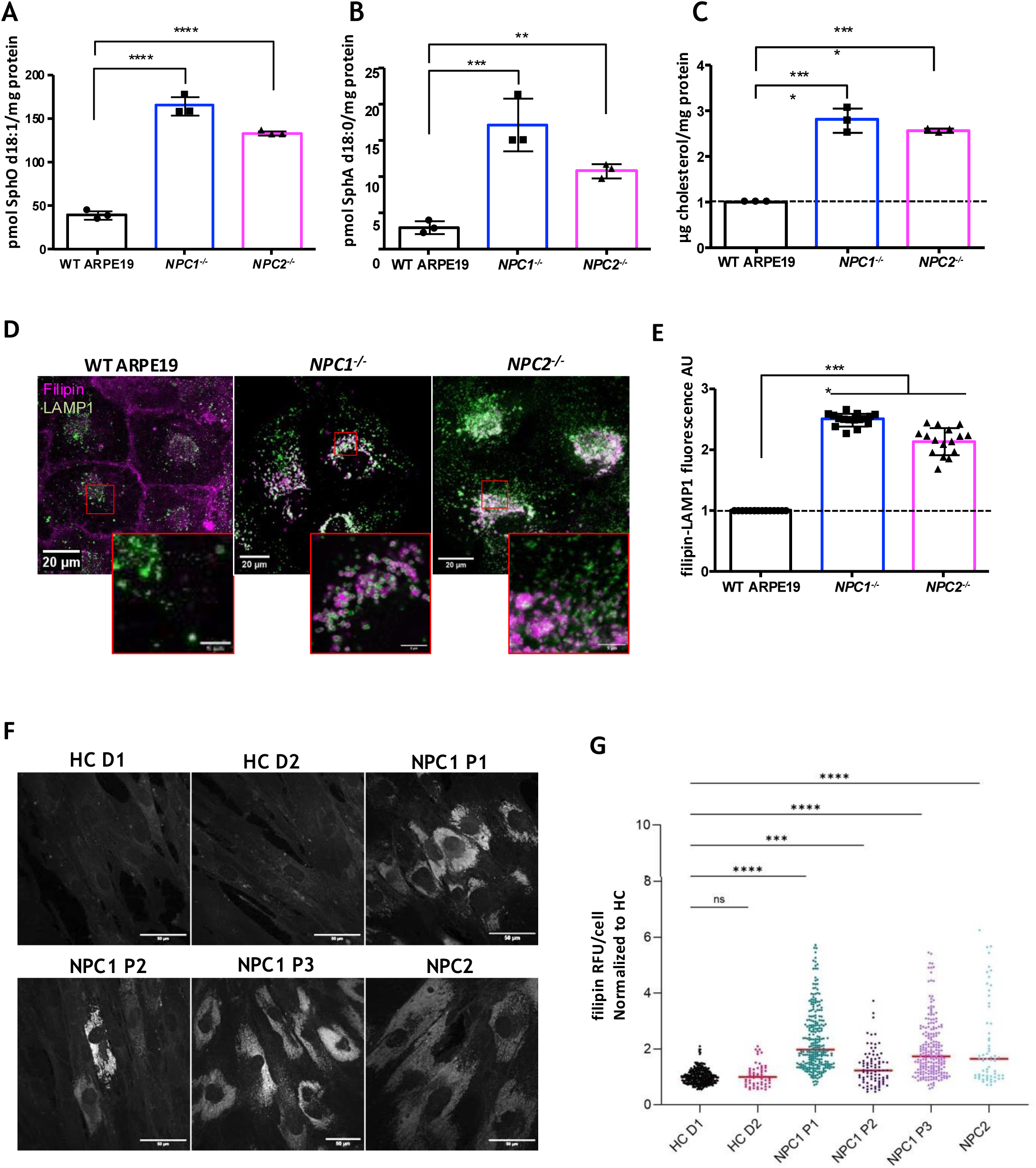
*NPC1^−/−^* and *NPC2^−/−^* ARPE-19 cells recapitulate classical NPC lipid accumulation patterns. **(A, B)** Quantification of **(A)** sphingosine (SphO d18:1) and **(B)** sphinganine (SphA d18:0) levels by HPLC in WT, *NPC1*⁻/⁻, and *NPC2*⁻/⁻ ARPE-19 cells. Levels are expressed as pmol per mg of total protein. Data shown are representative of 3 independent experiments. **(C)** Total cholesterol (esterified and unesterified) levels measured by Amplex Red assay in WT, *NPC1*⁻/⁻, and *NPC2*⁻/⁻ ARPE-19 cells. **(D)** Representative confocal microscopy images showing free cholesterol (filipin staining; pseudo-coloured magenta) co-localizing with LAMP1 (green). Scale bar: 5 μm. **(E)** High-content quantification of filipin signal intensity co-localizing with LAMP1-positive compartments (expressed in arbitrary units, AU). Each data point represents the average fluorescence derived from >1,000 cells. Data were normalized to WT ARPE-19 cells. Representative experiment of 3 biological replicates. **(F)** Filipin staining of primary human skin fibroblasts derived from NPC1 patients (P1–P3), an NPC2 patient (P1), and healthy control donors (HCD1, HCD2). Scale bar = 50μm. **(G)** Quantification of filipin signal intensity within the cytoplasm of HCDs and NPC patient fibroblasts. RFU/cell indicates the Relative Fluorescence Unit per cell. Statistical analysis was performed using one-way ANOVA followed by Dunnett’s multiple comparisons test. Data are presented as mean ± SD; ns = not significant; \*\*\*\**p* < 0.0001, \*\*\**p* < 0.001, \*\**p* < 0.01. Unless otherwise stated, n = 3 independent experiments.

Re-expression of NPC1 or NPC2 by lentiviral transduction reversed these NPC-like phenotypes, normalizing lysosomal morphology, reducing cholesterol accumulation, and decreasing LysoTracker intensity (Fig. 4). These results indicate that the observed defects actually arise from loss of NPC1 or NPC2, excluding any major Cas9 off target mutations that might occur because of sequence similarity (Kuscu et al., 2014).

**Figure 4:**
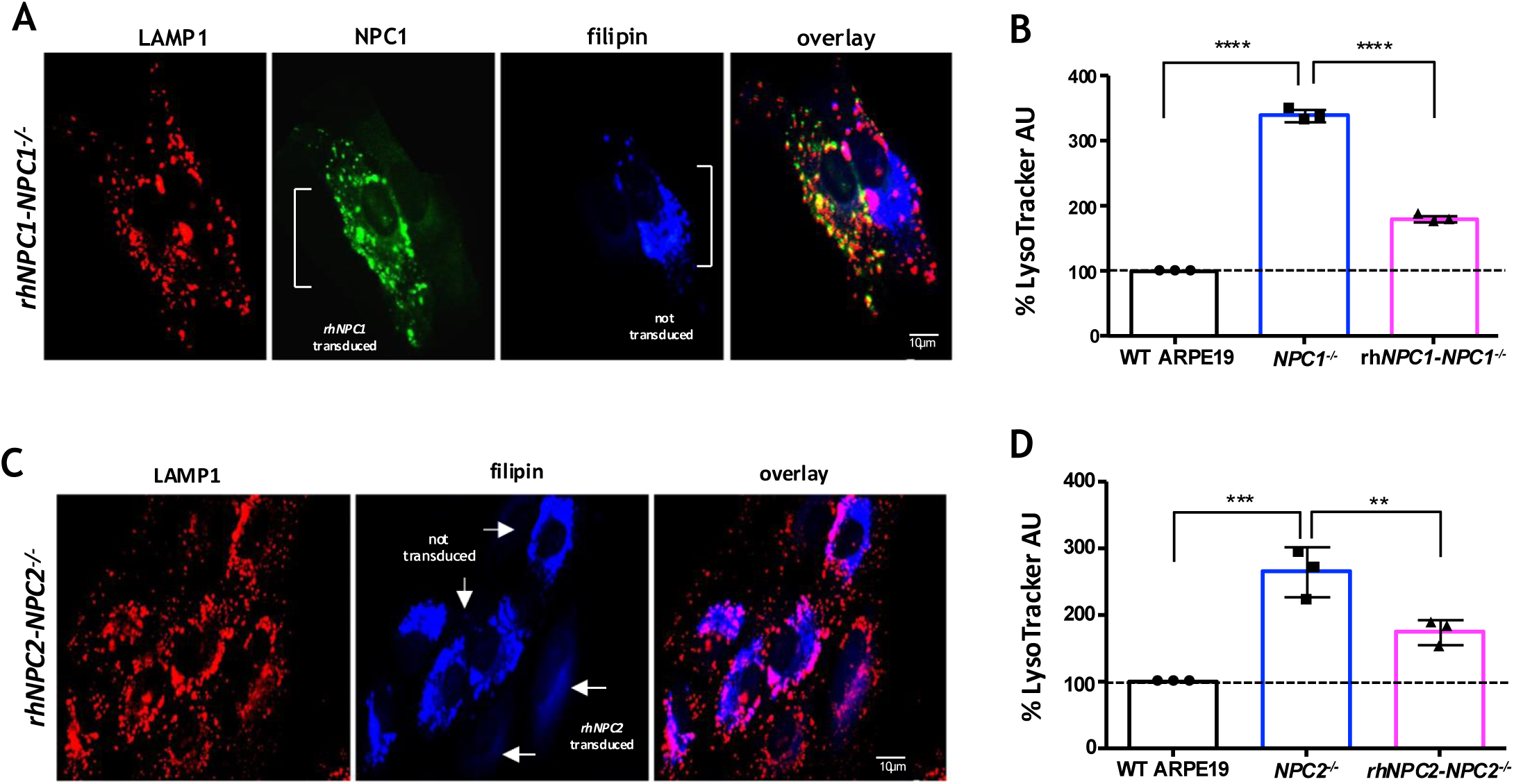
Reconstitution of NPC1 or NPC2 rescues lipid accumulation and lysosomal expansion. *NPC1⁻/⁻* or *NPC2⁻/⁻* ARPE-19 cells were transduced with lentiviral vectors encoding recombinant human NPC1 (rhNPC1) or NPC2 (rhNPC2), respectively. **(A)** Representative confocal microscopy images of a mixed population containing transduced (rhNPC1) and non-transduced *NPC1*⁻/⁻ ARPE-19 cells, stained for NPC1 (green), LAMP1 (red), and free cholesterol (filipin; blue). **(B)** Representative confocal microscopy images of a mixed population containing transduced (rhNPC2) and non-transduced *NPC2*⁻/⁻ ARPE-19 cells, stained for LAMP1 (red) and free cholesterol (filipin; blue). **(C, D)** Flow cytometric quantification of the percentage of LysoTracker Green-positive cells in WT, mixed transduced populations, and isogenic *NPC1*⁻/⁻ or *NPC2*⁻/⁻ ARPE-19 cell lines. Statistical analysis was performed using one-way ANOVA followed by Dunnett’s multiple comparisons test. Data are presented as mean ± SD from 3 independent experiments. \*\*\*\**p* < 0.0001; \*\**p* < 0.01.

### ARPE19 NPC knockout models enable multi-parametric high-content screening

The robust and uniform phenotypes of the ARPE19 *NPC1⁻^/^⁻* and *NPC2⁻^/^⁻* lines allow for the quantitative assessment of therapeutic candidates across multiple disease-relevant readouts. Co-staining with filipin and LAMP1 provided clear visualization of endolysosomal free cholesterol sequestration via both conventional and high-content confocal microscopy (Fig. 3D, 5A-C). This imaging profile, integrated with LysoTracker-based flow cytometry and HPLC quantification of sphingosine, establishes a multi-parametric platform for evaluating phenotypic reversal. To validate this approach, we utilized 2-hydroxypropyl-β-cyclodextrin as a reference compound (Rosenbaum et al., 2010). High-content analysis using a customized automated script demonstrated that β-cyclodextrin treatment normalized lysosomal cholesterol in *NPC2⁻^/^⁻* cells (Fig. 5A-D) and significantly reduced LysoTracker fluorescence in both *NPC1⁻^/^⁻* and *NPC2⁻^/^⁻* genotypes (Fig. 5F-G). Collectively, these data establish these isogenic models as sensitive, high-throughput *in vitro* platforms for the identification and validation of compounds targeting NPC-associated lipid dysregulation.

**Figure 5:**
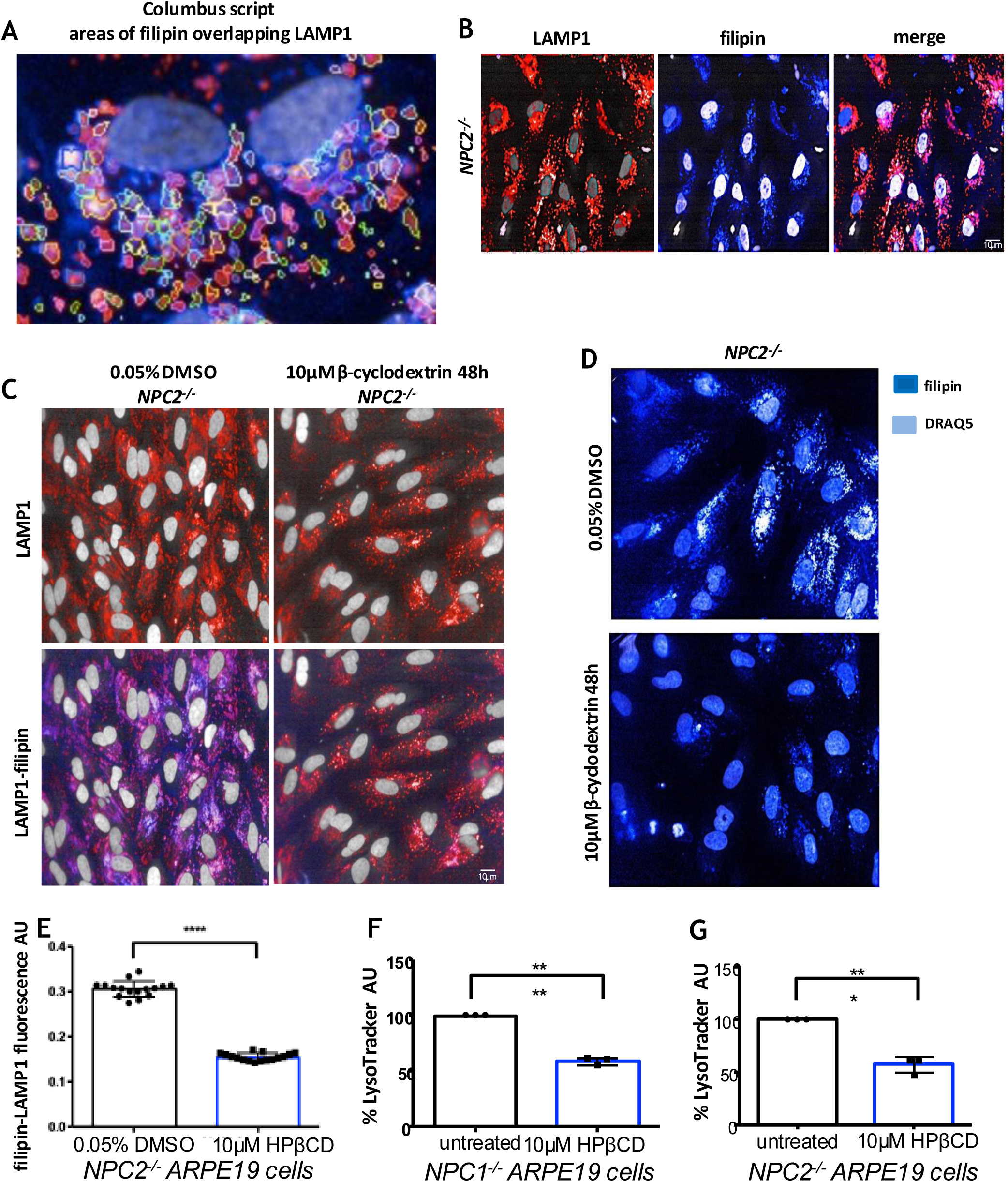
ARPE-19 *NPC1*⁻/⁻ and ARPE-19 *NPC2*⁻/⁻ cells as platforms for high-content drug screening. (A) Representative confocal microscopy images of LAMP1 and filipin staining used as a readout for lysosomal free cholesterol levels in *NPC1*⁻/⁻ and *NPC2*⁻/⁻ ARPE-19 cells. (B) Representative image of a Columbus script segmentation mask displaying the overlap between filipin and LAMP1 signals used to quantify lysosomal free cholesterol. (C) Representative images from Opera high-content screening (HCS) of ARPE-19 *NPC2*⁻/⁻ cells treated with 0.05% DMSO (control) or 10μM 2-hydroxypropyl-β-cyclodextrin (HPβCD) for 48 h. Cells were stained with LAMP1 (red), filipin (pseudo-coloured cyan) and DRAQ5 (nuclei, pseudo-coloured grey). (D) High-content quantification of filipin fluorescence intensity (expressed in arbitrary units, AU) co-localizing with LAMP1-positive compartments in *NPC2*⁻/⁻ ARPE-19 cells treated with DMSO or 10μM HPβCD for 48 h. Each data point represents the average fluorescence per cell calculated from >1,000 cells. (E) Representative images of ARPE-19 *NPC2*⁻/⁻ cells stained with filipin (pseudo-coloured cyan) and DRAQ5 (pseudo-coloured grey) showing clearance of accumulated free cholesterol following treatment with 10μM 2-hydroxypropyl-β-cyclodextrin (HPβCD) for 48 h. (F-G) Flow cytometric quantification of the percentage of LysoTracker Green-positive *NPC1*⁻/⁻ and *NPC2*⁻/⁻ ARPE-19 cells following treatment with DMSO or 10μM HPβCD for 48 h. Statistical analysis was performed using a two-tailed Student’s t-test. Data are presented as mean ± SD; **** *p* < 0.0001, *** *p* < 0.001. n = 3 independent experiments.

### Divergent effects of NPC1 and NPC2 deficiency on mitochondria-lysosome contact site remodeling

The downstream consequences of lysosomal lipid accumulation in NPC are complex and, perhaps surprisingly given the lysosomal primary defect, include mitochondrial dysfunction (Yu et al., 2005), (Ordonez et al., 2012) (Szilvia Kiraly et al., 2025). While the underlying mechanism of coupled lysosome and mitochondrial dysfunction in NPC has not yet been fully elucidated, we and others have identified expanded mitochondria:lysosome contact sites (MLCs) in NPC1 patient-derived and NPC1-inhibited cells (Höglinger et al., 2019), (Calì et al., 2025). MLC expansion in NPC is dependent on the LE/Lys lipid transport protein StARD3 (Höglinger et al., 2019) (S. Kiraly et al., 2025), previously shown to mediate mitochondrial cholesterol accumulation in NPC (Charman et al., 2010). Since excess mitochondrial cholesterol has been implicated in mitochondrial dysfunction, aberrant crosstalk at expanded MLCs may contribute to NPC disease pathogenesis. We therefore examined MLCs in *NPC1⁻/⁻*, and *NPC2⁻/⁻* ARPE19 cells using both fluorescence and electron microscopy. Proximity ligation assay (PLA) can provide a fluorescent read-out of MLCs (Fig. 6A) and using antibodies to LAMP1 on LE/Lys and TOM20 on mitochondria, identified an approximately two-fold increase in MLCs in cells lacking NPC1 (Fig. 6B and 6C), which was validated by electron microscopy analysis (Fig. 6D and 6E). In contrast, however, there was a modest reduction in MLCs in cells lacking NPC2 (Fig. 6B and 6C). These results are consistent with our previous reports in other cellular models of NPC (Höglinger et al., 2019) (Pastore et al., 2025) and with a role for expanded MLCs in cholesterol transport to the mitochondria, which is increased in cells lacking functional NPC1 but not in the absence of NPC2 (Kennedy et al., 2012)

**Figure 6:**
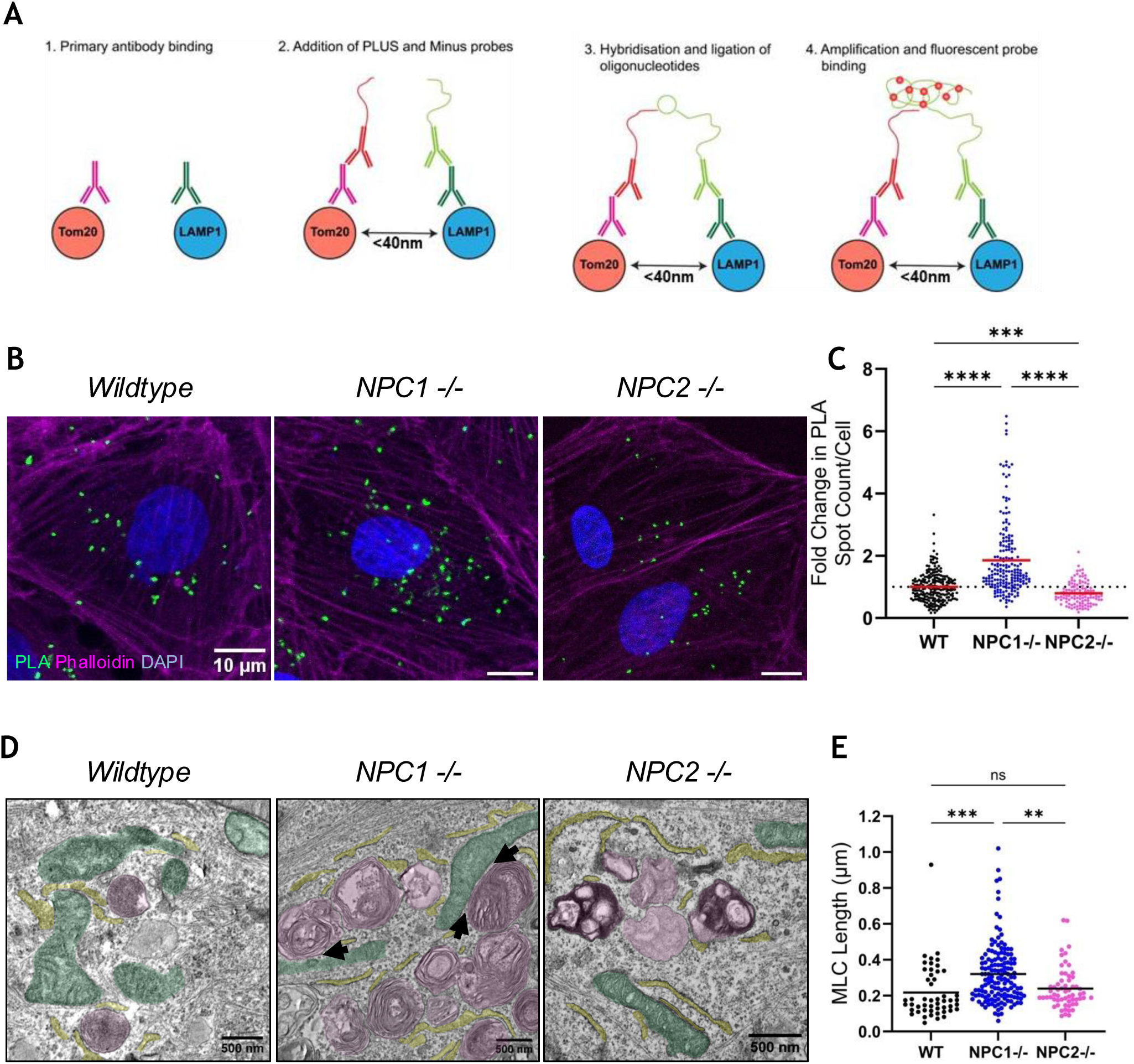
Expanded mitochondria-lysosome contact sites (MLCs) in *NPC1^−/−^* but not in *NPC2^−/−^* ARPE-19 cells. (A) Proximity ligation assay (PLA) schematic. (B) Representative images from the Mito-Lys PLA and (C) quantification of fold change in PLA spots per cell (normalised to wildtype) showing that MLCs are increased in ARPE-19 *NPC1^−/−^* cells and reduced in *NPC2^−/−^* cells compared to wildtype. Scale bars 10 µm. Statistical analysis performed using one-way ANOVA. (C) Representative TEM images showing LE/Lys (false-coloured magenta), mitochondria (false-coloured green) and ER (false-coloured yellow); black arrows indicate MLCs. (D) Quantification of MLC length (µm) from TEM images. Each point represents one MLC. Bar represents mean, statistical analysis performed using ordinary one-way ANOVA (n = 3 biological repeats).

### StARD3-targeted APEX2 proximity labelling identifies HKDC1 as a mitochondrial regulator of MLCs in NPC

In addition to our previous finding that MLC expansion in NPC1 patient fibroblasts or NPC1-inhibited HeLa cells is StARD3-dependent (Höglinger et al., 2019); (S. Kiraly et al., 2025), STARD3 was also shown by electron tomography to form visible tethers at MLCs (Nara et al., 2023). As expected, StARD3-GFP was localised at LAMP1-positive LE/Lys in *NPC1^⁻/⁻^* ARPE19 (Fig. 7A), confirmed by co-localisation analysis (Fig. 7B). Consistent with a role for StARD3 in MLC tethering in ARPE19 cells, PLA revealed a significant increase in MLCs in WT ARPE19 cells overexpressing StARD3, with a small (not significant) additional increase in MLC numbers on StARD3 overexpression in *NPC1^⁻/⁻^* ARPE19 cells (Fig. 7C). While StARD3 has previously been shown to regulate MLCs in NPC, the mitochondrial tethering partner in non-steroidogenic cells has not yet been described (Y. Xie et al., 2024), (Yinyin Xie et al., 2024).

**Figure 7:**
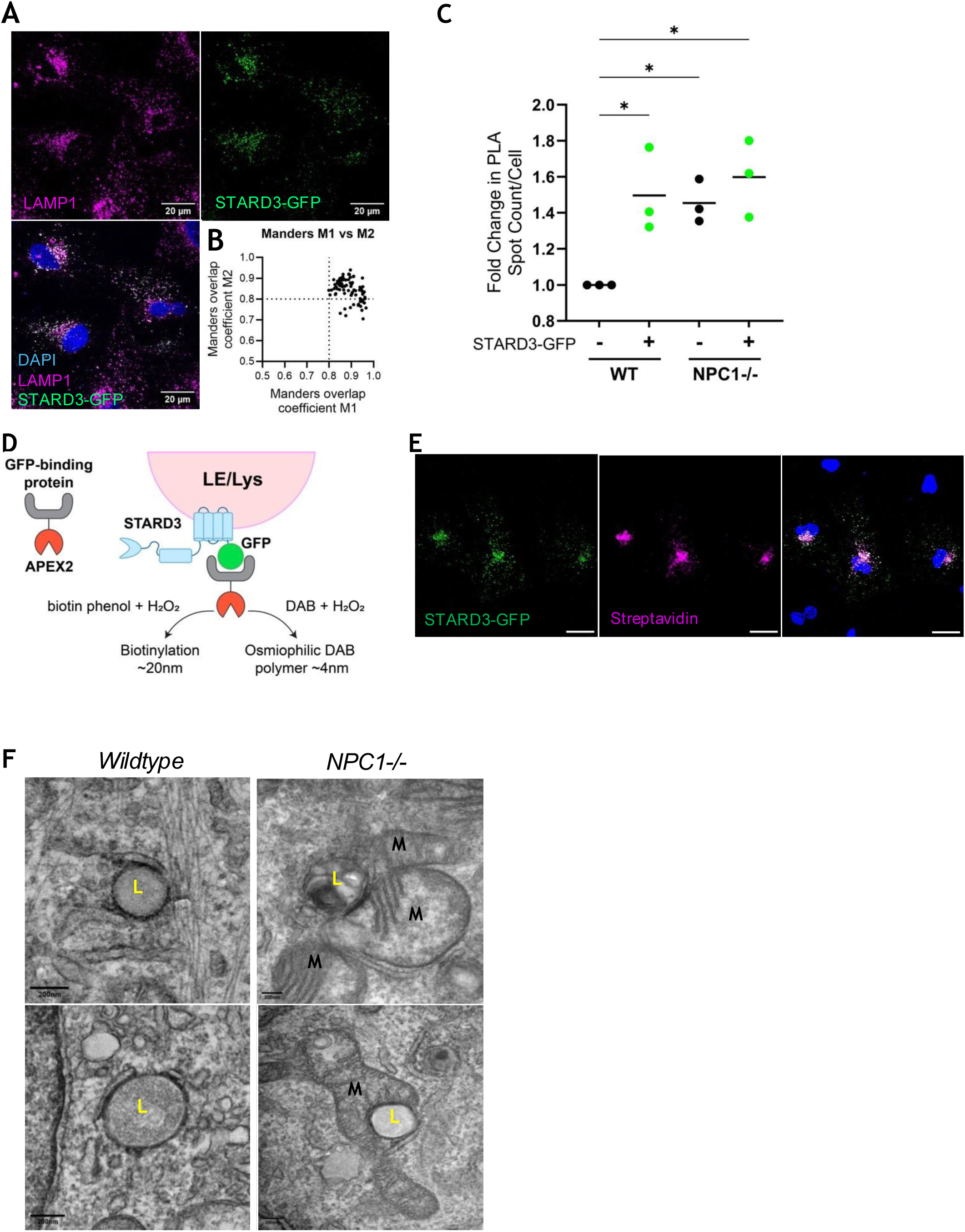
APEX targeting to expanded MLCs in NPC using StARD3-GFP. A) Cells expressing StARD3-GFP were stained for the LE/Lys marker LAMP1 and imaged by confocal microscopy. B) Co-localisation of endogenous LAMP1 and over-expressed STARD3-GFP, quantified using Manders’ Colocalisation, confirming localisation of STARD3-GFP at LAMP1-positive LE/Lys. C) Ly-Mt PLA STARD3 in ARPE-19 cells ± StARD3 GFP overexpression. Each point represents mean fold change in PLA spots normalised to wildtype non-transfected control. N = 3 biological repeats, minimum 20 cells per experiment. D) Schematic of STARD3-GFP/APEX2-GFP-Binding Peptide (csGBP)-mediated biotin labelling. E) Validation of csGBP-APEX2 localisation by IF and F) by TEM following DAB staining.

To identify StARD3 interactions at the expanded MLCs in *NPC1^⁻/⁻^* ARPE19 cells (Fig. 6), a STARD3-targeted proximity labelling approach was taken, using an engineered ascorbate peroxidase enzyme, APEX2, conjugated to a conditionally stable GFP-binding peptide (csGBP) to biotinylate proteins within an approximately 20 nm radius of StARD3-GFP (Fig. 7D). Labelling biotin with fluorescent streptavidin confirmed biotinylation of proteins in close proximity to StARD3 in *NPC1^⁻/⁻^* ARPE19 cells co-expressing StARD3-GFP and APEX2-csGBP (Fig. 7E). APEX2 also catalyses the polymerisation of diaminobenzidine (DAB) to create an electron-dense reaction product that can be visualised by electron microscopy (Tan et al., 2020). Using this method, StARD3-GFP/APEX2-csGBP localisation to LE/Lys was confirmed in both wild-type and *NPC1^⁻/⁻^* ARPE19 cells. Whereas in wild-type StARD3-GFP/APEX2-csGBP-expressing cells the electron-dense DAB reaction product decorated the ER:LE/Lys interface, in *NPC1^⁻/⁻^* ARPE19 cells, it was enriched at MLCs (Fig. 7F). Together these data demonstrate APEX2-csGBP localisation to and activity at StARD3-regulated contact sites.

StARD3 proximity labelling was compared in wild-type and NPC1-knockout ARPE19 cells by mass spectrometry (MS). Direct comparisons of label-free quantification (LFQ) intensities of proteins, revealed 124 differentially biotinylated proteins in *NPC1^−/−^* compared with wild-type ARPE19 cells (> 2-fold change), with a *p*-value <0.05 (Fig. 8A). Of these, only four proteins were mitochondrial, with hexokinase domain containing protein 1 (HKDC1) the most significantly differentially biotinylated (Fig. 8A). HKDC1 has previously been implicated in MLC regulation (Cui et al., 2024) and was therefore a strong candidate as a component of an MLC tethering complex that is upregulated in NPC. Western blotting of wild-type and ARPE19 *NPC1^−/−^* cells revealed a marked increase in HKDC1 expression in cells lacking NPC1 (Fig. 8B and 8C) and overexpression of HKDC1-GFP in wild-type ARPE19 cells resulted in a modest but significant increase in MLC numbers, consistent with a role for HKDC1 in tethering StARD3-dependent MLCs in NPC.

**Figure 8:**
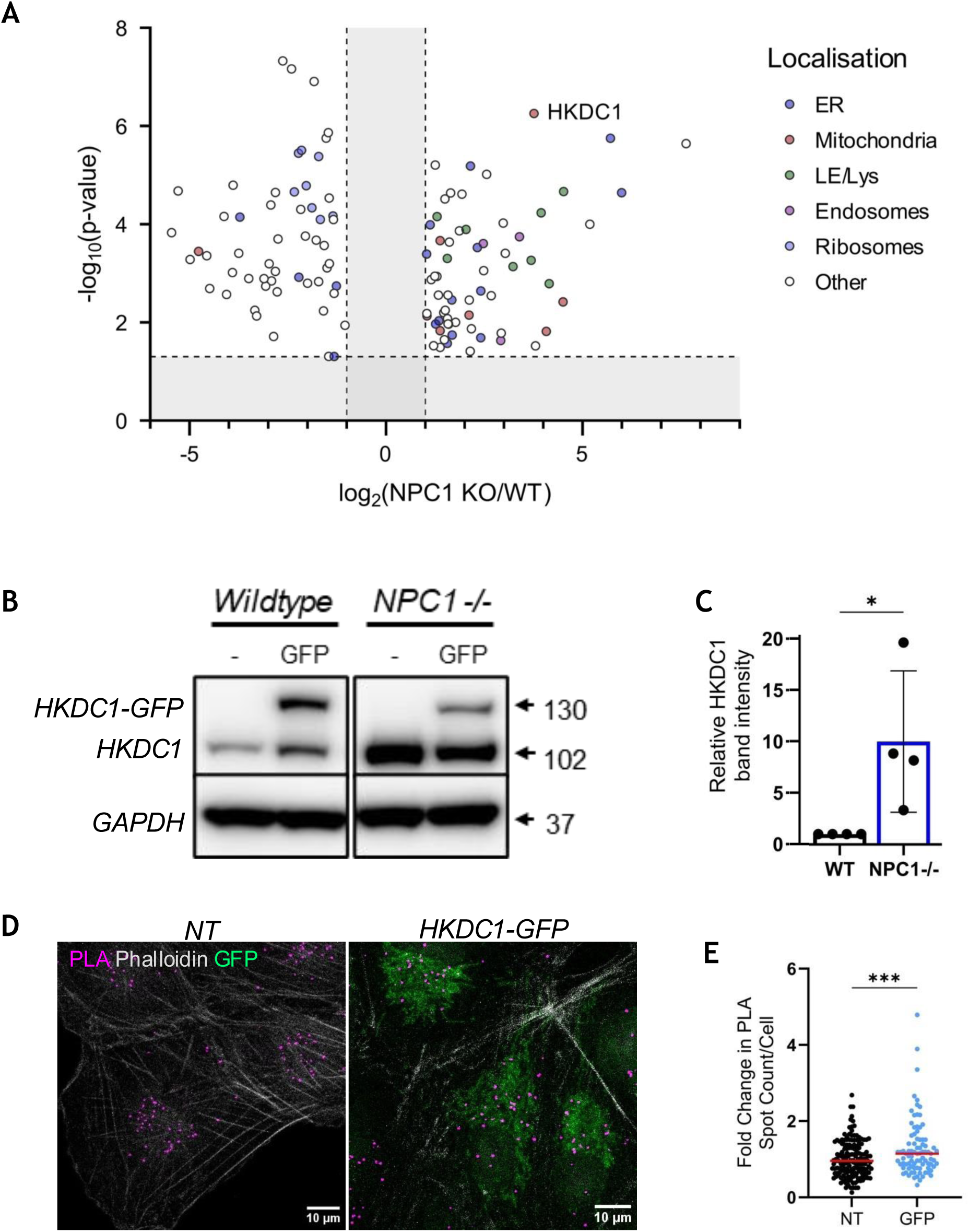
STARD3-targeted APEX2-mediated proximity labelling screen identified HKDC1 as a novel MLC regulator in ARPE-19 *NPC1^−/−^* cells. A) Volcano plots presenting biotinylated proteins identified by mass spectrometry. The x axis of volcano plots specifies the logarithm to the base 2 of fold change (FC) in label-free quantification (LFQ) intensities, and the y axis specifies the negative logarithm to the base 10 of the t test p-values. Fold change (log2FC) displayed in volcano plots directly comparing protein biotinylation in wildtype ARPE-19 cells with *NPC1^−/−^* cells. Proteins colour-coded by subcellular localisation. Vertical dashed lines at log2(FC) = 1 (FC = 2) represent fold change cut off, horizontal lines at -log10(P-value) = 1.301 (p = 0.05). B) Immunoblotting identified HKDC1 is overexpressed in *NPC1^−/−^* ARPE-19 cells and confirmed overexpression of HKDC1-GFP. C) HKDC1 band intensity normalised to GAPDH loading control. Bar represents mean with SD, each point represents biological repeat. Statistical analysis performed using unpaired t test. D) Representative images of Mito-Lys PLA in non-transfected or HKC1-GFP overexpressing wildtype ARPE-19 cells. E) Quantification of PLA spot count per cell identified HKDC1-GFP overexpression increased MLCs. Each point represents one cell, N = 3, minimum 113 cells over 3 experiments. Bar represents mean. Statistical analysis performed using unpaired t test.

## Discussion

Patients with loss of NPC1 or NPC2 are clinically indistinguishable (Vanier, 2010), but differences in cellular phenotypes are starting to emerge, particularly in the contact site landscape. We have developed human isogenic models of NPC1 and NPC2 deficiency that recapitulate lysosomal storage disease phenotypes observed in patient-derived cells, including homogeneous lysosomal expansion and lipid sequestration (Fig. 2C, 3A, 4). Notably, while patient-derived fibroblasts exhibit surprising variability in lysosomal size and cholesterol levels both within and between cell lines (Fig. 3F), the ARPE19 *NPC1^−/−^* and *NPC2^−/−^* cells provide a highly consistent phenotypic readout (Fig. 2D). This suggests that the phenotypic inconsistency observed in primary lines is likely influenced by mutation-specific effects or patient-specific genetic variables, highlighting the importance of isogenic systems for direct comparison.

We previously identified variation in the extent of ER interaction with LE/Lys in cells from NPC1 and NPC2 patients. Using a combination of fluorescence and electron microscopy (Höglinger et al., 2019) as well as proximity ligation assays (Szilvia Kiraly et al., 2025), we found reduced ER:LE/Lys contact in cells from NPC1 but not NPC2 patients and demonstrated NPC1 enrichment at ER contact sites, where it associates with the ER-localised lipid transfer protein Gramd1b (Höglinger et al., 2019). Whether NPC1 directly interacts with Gramd1b, or generates cholesterol-rich microdomains on the LE/Lys membrane that result in Gramd1b recruitment as has been shown following cholesterol enrichment at other membranes (Sandhu et al., 2018), has not yet been established. In contrast to findings in the patient-derived cells used in our study, ER contact with LE/Lys has also been reported to be increased in NPC1-deficient cells (Lim et al., 2019), likely reflecting underlying variation between the patient cell lines used.

As well as variance in contact site extent between the ER and LE/Lys in cells from NPC1 compared with NPC2 patients, there are also pronounced differences in mitochondrial contact with LE/Lys. In contrast to the expanded MLCs in NPC1-inhibited cells or NPC1 patient-derived fibroblasts (Höglinger et al., 2019) (Giamogante et al., 2024) (S. Kiraly et al., 2025), MLCs were reduced in HEK cells lacking NPC2 (Pastore et al., 2025). However, the contribution of the different genetic backgrounds to these divergent phenotypes is not known, highlighting the need for isogenic models of NPC1 and NPC2 to enable direct comparisons. Our data here confirm that MLCs are increased in cells lacking NPC1 compared with NPC2. We have previously identified a correlation between lysosomal cholesterol and MLC extent in primary fibroblasts from NPC1 patients (S. Kiraly et al., 2025). The slight reduction in MLCs in cells lacking NPC2, where LE/Lys cholesterol accumulation is comparable with that in *NPC1* knockout cells (Fig. 3B-D), suggests that accessibility of the LE/Lys cholesterol stores may be a key factor in MLC regulation. How cholesterol is transported to the LE/Lys limiting membrane in the absence of NPC1 is not completely understood, but in addition to NPC1, two other lysosomal membrane proteins, LIMP2 (Heybrock et al., 2019) and LAMP2 (Schneede et al., 2011) have been implicated in the egress of LDL-derived cholesterol from LE/Lys. While NPC2 has been suggested to function in the cholesterol binding function of LAMP2, there is no endogenous interaction detected between LIMP2 and NPC2, suggesting they may function separately (Heybrock et al., 2019). Interestingly a role for LIMP2 at ER:LE/Lys contact sites has been reported, with lysosomal LIMP2 in a complex with ER-localised VAP through StARD3 interaction (Rudnik et al., 2024).

The expansion of MLCs in NPC1- but not NPC2-deficient cells, is consistent with a potential role for these contacts as conduits for cholesterol transport to mitochondria, with StARD3-dependent mitochondrial cholesterol accumulation reported in cells lacking functional NPC1 but not NPC2 (Kennedy et al., 2012). Mitochondrial cholesterol accumulation is strongly implicated in oxidative stress and changes in metabolic homeostasis (Kennedy et al., 2014) (Balboa et al., 2017), and is therefore likely to contribute to mitochondrial dysfunction in NPC. Interesting differences in mitochondrial phenotypes have been reported in cellular models of NPC1 and NPC2 disease. In contrast to the widely described defect in mitochondrial respiration and reduced mitochondrial membrane potential in cells lacking functional NPC1 (Yu et al., 2005) (Ordonez et al., 2012) (Szilvia Kiraly et al., 2025), mitochondrial respiration was unperturbed in NPC2-deficient HEK cells, while mitochondrial membrane potential was increased, suggesting a subtle hyper-polarisation phenotype (Pastore et al., 2025). Together these findings implicate MLC expansion and consequent cholesterol import in mitochondrial dysfunction.

The shared genetic background, ease of use (including rapid growth and ready transfection), and excellent phenotypic consistency lend the ARPE19 cell lines well to high-throughput screening. We found that, as has been reported for other models of NPC, LysoTracker intensity (reflecting lysosomal expansion) and accumulation of filipin-stained cholesterol in LAMP1-positive LE/Lys (Fig. 3C-D) provide reliable readouts of NPC disease phenotypes in both ARPE19 *NPC1^−/−^* and ARPE19 *NPC2^−/−^* cells. Our data showing that treatment with β-cyclodextrin significantly reduced the intensity of both lysotracker signal and filipin-stained cholesterol in the LAMP1-positive compartment in ARPE19 *NPC2^⁻/⁻^* cells (Fig. 5D-E) demonstrate the suitability of ARPE19 cellular models of NPC for high content screening for compounds that can reverse NPC phenotypes. Proximity labelling is another commonly used screening tool and using StARD3 as a known MLC regulator (Höglinger et al., 2019) (Szilvia Kiraly et al., 2025; S. Kiraly et al., 2025), we conducted a StARD3-targeted APEX2-mediated proximity labelling screen in ARPE19 *NPC1^−/−^* compared with WT cells. Analysis of biotinylated proteins by mass spectrometry identified increased proximity of the mitochondrial protein HKDC1 to StARD3 in the absence of NPC1. HKDC1 has been previously implicated at MLCs (Cui et al., 2024), making it a strong candidate tethering complex component for the expanded MLCs in NPC1-deficient cells. While HKDC1 is characterized as a hexokinase (Piwko et al., 2025), its role in regulating MLCs appears to be independent of its catalytic activity, functioning instead as a structural scaffold. Recent evidence indicates that under lysosomal stress, HKDC1 is recruited to the outer mitochondrial membrane (OMM) where it interacts with the TOM complex components, TOM20 and TOM70 (Cui et al., 2024), as well as with VDAC2 (Voltage-Dependent Anion Channel 2), facilitating the formation of MLCs, a process regulated via the VDAC2/Ras-PI3K axis (Satoh et al., 2023). Loss of HKDC1 was shown to result in a significant reduction in mitochondria-lysosome contact, affecting lysosomal homeostasis (Cui et al., 2024). In our model, the 10-fold upregulation of HKDC1 likely contributes to the observed expansion of MLCs. MLC expansion may serve a critical role in metabolic buffering, likely providing platforms for the direct transfer of lipids, including StARD3-dependent cholesterol transport (Wilhelm et al., 2017) or contact-dependent signaling to mitigate the consequences of NPC1 deficiency (Charman et al., 2010; Kennedy et al., 2012). Interestingly VDAC1 interaction with StARD3 was recently reported in aldosterone-producing adenomas, facilitating StARD3-dependent mitochondrial cholesterol import for aldosterone production (Chen et al., 2025); a StARD3-HKDC1-VDAC axis may prove important for the mitochondrial cholesterol accumulation and dysfunction in NPC.

The significant upregulation of HKDC1 that we observed in ARPE19 *NPC1⁻/⁻* cells identifies a novel mechanism coupling lysosomal stress with mitochondrial function involving organelle contact site remodelling. Increased HKDC1 expression could, at least in part, account for the increase in HKDC1 biotinylation in ARPE19 *NPC1^⁻/⁻^* cells. However, our finding that MLC numbers are increased in wild-type cells overexpressing HKDC1, together with previous reports of reduced MLCs in HKDC1-depleted cells (Cui et al., 2024), strongly supports a role for HKDC1 in MLC regulation. HKDC1 was recently identified as a direct transcriptional target of TFEB, specifically upregulated in response to both mitochondrial and lysosomal dysfunction (Cui et al., 2024). In our model, the accumulation of lysosomal lipids likely triggers the nuclear translocation of TFEB as part of a coordinated response to lysosomal dysfunction. Increased TFEB activity has been reported in several NPC models (Willett et al., 2017) (Contreras et al., 2020), (Davis et al., 2025), and in ARPE19 *NPC1^−/−^* cells, expression of LAMP1, which is regulated by TFEB, was 3-fold higher than in wildtype ARPE19 cells, consistent with increased TFEB activity (Supplementary Fig. S2A and S2B). TFEB activation typically occurs when lysosomal stress leads to the inhibition of mTORC1 (mechanistic Target of Rapamycin Complex 1) or the TRPML1 (MCOLN1)-mediated activation of calcineurin, both of which promote TFEB dephosphorylation and subsequent nuclear translocation (Settembre et al., 2012), (Medina et al., 2015). On the other hand, TFEB phosphorylation by active mTORC1 prevents its nuclear translocation, down-regulating transcriptional activity. Since hyperactivation of mTORC1 was previously demonstrated in NPC1-deficient cells (Davis et al., 2021), mTORC1-independent mechanisms (Schwendener Forkel et al., 2024) must drive the increased TFEB activity in NPC. Similarly, defects in lysosomal Ca^2+^ stores and reduced release through the LE/Lys Ca^2+^ channel TRPML1 have been reported in NPC (Lloyd-Evans et al., 2008) (Shen et al., 2012), suggesting that TFEB activation in NPC is likely to be independent of calcineurin as well as mTORC1. Additional regulators of TFEB activation have been reported, including direct phosphorylation by MAPK (Martina et al., 2022) as well as TFEB acetylation, which promotes nuclear translocation and lysosome biogenesis (Li et al., 2022), but the precise mechanism of increased TFEB activity in NPC remains unclear.

In summary, the isogenic NPC1 and NPC2 deficient ARPE19 cells generated here reflect the lysosomal storage disease phenotypes seen in patient-derived cells but with improved phenotypic consistency and without any potentially complicating effects of different genetic backgrounds. MLC analysis in these cells demonstrated a marked difference in inter-organelle interaction in isogenic models of NPC1 and NPC2 deficiency, revealing that NPC2-dependent cholesterol transport is required for the previously observed correlation between LE/Lys cholesterol accumulation and MLC expansion in cells lacking functional NPC1. Through our proximity labelling screen we have identified HKDC1 as a mitochondrial protein that is overexpressed in ARPE19 *NPC1^−/−^* and contributes to MLC expansion, suggesting a potential role for HKDC1 in the pathogenesis of NPC.

## Materials and Methods

### Cell culture

ARPE-19 cells (ATCC® CRL-2302) were maintained in DMEM/F12 supplemented with 10% fetal bovine serum (FBS), 1% penicillin/streptomycin, and 1% L-glutamine at 37 °C with 5% CO₂.

Primary fibroblasts derived from NPC1 and NPC2 patients carrying disease-causing mutations and fibroblasts from healthy donors were obtained from the Corriell Institute (HC1: GM05757, HC2: GM05399, NPC1-P1: GM18398, NPC1-P2: GM17921, NPC1-P3: GM22871).

### Generation of *NPC1⁻/⁻* and *NPC2⁻/⁻* ARPE-19 cell lines

CRISPR/Cas9-mediated knockout was performed using the “All-in-One” GFP-Cas9 plasmid (Sigma-Aldrich). For NPC1 editing, a single guide RNA (sgRNA) targeting exon 1 (5ʹ-GCTGCTACTGTGTCCAGCGC-3ʹ) was used; for NPC2, an sgRNA targeting exon 3 (5ʹ-CTGATGGTTGTAAGAGTGGA-3ʹ) was employed. ARPE-19 cells (1 × 10⁶) were transfected by nucleofection (Lonza, VCA-1003). After 72 h, GFP⁺ cells were single-cell sorted by fluorescence-activated cell sorting (FACS) into 96-well plates. Genomic DNA from expanded clones was analyzed by Sanger sequencing. A clone carrying a homozygous c.56delC mutation in NPC1 and a clone with a homozygous c.226delG mutation in NPC2 were selected for further studies.

### Filipin staining

Unesterified cholesterol was visualized by filipin III staining (Abcam). Cells were fixed in 4% paraformaldehyde, quenched with 50 mM glycine, and incubated with filipin III (diluted 1:100 in kit solvent) for 1 h at room temperature in the dark. Imaging was performed on a confocal microscope (excitation 340–380 nm, emission 385–470 nm).

### Sphingosine and sphinganine quantification by HPLC

Sphingoid bases were extracted from cell lysates, processed and analyzed as previously described (Merrill et al., 2000), (Danielle te Vruchte, 2025). Briefly, samples containing 0.1 mg protein were spiked with C20-sphingosine internal standard, subjected to solid-phase extraction on NH₂ columns (Biotage), and eluted with acetone. Dried extracts were resuspended in ethanol, derivatized with O-phthaldialdehyde (OPA), and analyzed by reverse-phase HPLC using a C18 column (Waters). Detection was performed with a fluorescence detector (excitation 340 nm, emission 455 nm).

### BODIPY-LacCer trafficking

WT, *NPC1^−/−^*, and *NPC2^−/−^* ARPE19 cells were pulsed with 10μM BODIPY-LacCer/BSA complexes for 15 min at room temperature and subsequently washed and chased in medium containing 1% FBS for 45min at 37°C (Chen et al., 1999). Live-cell fluorescence microscopy was performed using a Leica SP8 confocal microscope with identical acquisition settings across all genotypes. Images shown are representative of hundreds of cells examined across more than three independent experiments. Image assessment was performed by an external observer blinded to cell line identity.

### LysoTracker assay and flow cytometry

Lysosomal content was quantified using LysoTracker Green DND-26 (Thermo Fisher). ARPE-19 cells were stained with 200 nM LysoTracker Green in FACS buffer (0.1% BSA, 0.02 M NaN₃ in PBS) for 10 min at room temperature. Propidium iodide (1 µg/mL) was added immediately before acquisition to exclude dead cells. Data were collected on a BD FACS Canto II and analyzed with FACSDiva software (BD Biosciences) from at least 10,000 singlet events per condition.

### Western blotting

Cells were lysed in Laemmli buffer, and 30 µg of protein were resolved on 4–12% polyacrylamide gels (NuPAGE Novex). Proteins were transferred to PVDF membranes (Bio-Rad), blocked in 5% milk/PBS-T, and incubated overnight at 4 °C with primary antibodies: NPC1 (ab134113, Abcam, 1:200), NPC2 (HPA000835, Atlas, 1:200), LAMP1 (1D4B, DSHB, 1:500), LC3B (Cell Signaling 3868, 1:200). HRP-conjugated secondary antibodies were applied for 1.5 h at room temperature. Blots were developed with ECL (Amersham RPN2106) and imaged using a Bio-Rad ChemiDoc XRS+. Band intensities were quantified using Fiji.

### Cholesterol measurement

Cellular cholesterol was extracted following the Folch method and quantified using the Amplex Red Cholesterol Assay Kit (Molecular Probes) according to the manufacturer’s protocol.

### Immunofluorescence and High-Content Imaging

ARPE19 cells were seeded on glass coverslips or black-walled, clear-bottom 96-well plates and fixed with 4% paraformaldehyde (PFA) for 10 min at room temperature (RT). Cells were subsequently permeabilized and blocked for 45 min in PBS containing 2% bovine serum albumin (BSA) and 0.1% saponin. Primary antibodies were diluted in blocking buffer and incubated overnight at 4°C: mouse anti-LAMP1 (H4A3; Abcam, ab25630; 1:500), or rabbit anti-NPC1 (EPR5209; Abcam, ab134113; 1:200). After three 5-min washes in 1X PBS, cells were incubated for 1h at RT with Alexa-Fluor-conjugated secondary antibodies (goat anti-mouse-594 or goat anti-rabbit-488; 1:1000). Coverslips were mounted using VECTASHIELD® mounting medium. For subcellular localization analysis, confocal images were acquired using a Leica SP8 confocal microscope equipped with a 63x/1.4 NA oil-immersion objective and processed using Fiji/ImageJ. For high-content imaging (HCI) experiments, images were acquired using an Operetta High-Content Imaging System (PerkinElmer) and automated data analysis was performed using Columbus software (PerkinElmer).

### Proximity Ligation Assay (PLA)

The Duolink In Situ Detection Reagents Red (Sigma, DUO92008) kit was used for PLA, according to manufacturer’s instructions, with primary antibodies targeting LE/Lys (rabbit anti-LAMP1, CST #9091, 1:1000) and mitochondria (mouse anti-TOM20, sc-17764, 1:1000).

### Transmission Electron Microscopy (TEM)

Samples were prepared for TEM as previously described (Barral et al., 2022). Briefly, cells were fixed with 2% paraformaldehyde/ 2% glutaraldehyde in 0.1M sodium cacodylate buffer and post-fixed in 1.5% potassium ferricyanide/1% osmium tetroxide. Following treatment with UA-Zero (Agar Scientific, UK), cells were dehydrated through an ethanol series prior to embedding in resin.

Ultrathin sections were imaged on a JEOL 1400Plus microscope (JEOL ltd, Tokyo, Japan) fitted with an Advanced Microscopy Technologies (AMT) NanoSprint12 camera (AMT Imaging Direct, Woburn, MA, USA). Images were analysed using ImageJ software to measure parameters of lysosomes outlined manually using the freeform drawing tool. GraphPad Prism software was used to generate charts and non-parametric one-way ANOVA Kruskal-Wallis analysis.

### APEX2-mediated Proximity Labelling Screen

Cells transfected with STARD3-GFP (a kind gift from Fabien Alpy) and mKate2-P2A-APEX2-csGBP (a kind gift from Rob Parton (Addgene plasmid # 108875)) were treated with 2.5 mM biotin phenol for 30 minutes to increase bioavailability of biotin prior to addition of a reaction solution containing 1 mM H_2_O_2_ for one minute. The reaction was quenched with 20 mM sodium ascorbate, 10 mM sodium azide, and 5 mM Trolox in PBS.

*For immunofluorescence*: Cells were fixed with 4% PFA and biotin labelled using fluorescent streptavidin-647.

*For TEM*: Cells were fixed for TEM as detailed above and fixed cells were incubated with 3,3’-diaminobenzidine (DAB) solution containing 0.075% DAB (TAAB, D040), 10 mM imidazole (Sigma-Aldrich, I0250), 0.01% H_2_O_2_ in 0.1M Tris-HCl for 45min prior to postfix with osmium and processing for conventional TEM as described above.

*For mass spectrometry*: Biotinylated proteins were precipitated from cell lysates using magnetic streptavidin beads (MagResyn Streptavidin MS, MR-STP002). Following an overnight incubation with lysate at 4°C, beads were washed with 0.5 M KCl and 25 mM ammonium bicarbonate (Merck, 5330050050) prior to snap-freezing and storage at −80°C.

#### Mass Spectrometry

Bead-bound proteins were prepared for mass spectrometric analysis by in solution enzymatic digestion. Briefly, bead-bound proteins in 50 ul of 50 mM NH4HCO3 were reduced in 10 mM DTT, and then alkylated with 55 mM iodoacetamide. After alkylation, 0.5 ug of Trypsin (Promega, UK) was added and the proteins digested for 1 h at 37 °C in a thermomixer (Eppendorf, Germany), shaking at 800 rpm. Following this initial digestion, a further 0.5 ug of Trypsin (ThermFisherScientific, USA) was added and digestion continued overnight at 37 °C. The resulting peptides were analysed by nano-scale capillary LC-MS/MS using an Ultimate U3000 HPLC (ThermoScientific Dionex, San Jose, USA) to deliver a flow of approximately 250 nL/min. A C18 Acclaim PepMap100 5 µm, 75 µm x 20 mm nanoViper (ThermoScientific Dionex, San Jose, USA), trapped the peptides prior to separation on a C18 Acclaim PepMap RSLC 3 µm, 75 µm x 500 mm nanoViper (ThermoScientific Dionex, San Jose, USA). Peptides were eluted with a 90 min gradient of acetonitrile (2%v/v to 80%v/v). The analytical column outlet was directly interfaced via a nano-flow electrospray ionisation source, with a hybrid dual pressure linear ion trap mass spectrometer (Orbitrap Eclipse, ThermoScientific, San Jose, USA). Data dependent analysis was carried out, using a resolution of 120,000 for the full MS spectrum, followed by ten MS/MS spectra in the linear ion trap. MS spectra were collected over a m/z range of 275–1500. MS/MS scans were collected with the standard AGC target, dynamic maximum injection time mode, isolation window at 1.2 m/z and 32% normalised HCD collision energy. All raw files were processed with MaxQuant 2.4.9.0 (Cox & Mann, 2008) using standard settings and searched against a UniProt Human Reviewed KB database with the Andromeda search engine (Cox et al., 2011) integrated into the MaxQuant software suite. Enzyme search specificity was Trypsin/P for both endoproteinases. Up to two missed cleavages for each peptide were allowed. Carbamidomethylation of cysteines was set as fixed modification with oxidized methionine and protein N-acetylation considered as variable modifications. The search was performed with an initial mass tolerance of 6 ppm for the precursor ion and 0.5 Da for MS/MS spectra. The false discovery rate was fixed at 1% at the peptide and protein level. Statistical analysis was carried out using the Perseus module (v2.0.11) of MaxQuant. Prior to statistical analysis, peptides mapped to known contaminants, reverse hits and protein groups only identified by site were removed. Only protein groups identified with at least two peptides, one of which was unique and two quantitation events were considered for data analysis using Perseus software (Tyanova et al., 2016).

### Lentiviral Transduction of NPC1 and NPC2

To restore protein expression, *NPC1*⁻/⁻ and *NPC2*⁻/⁻ ARPE-19 cells were transduced with lentiviral vectors encoding human *NPC1* or *NPC2* (kindly provided by the Platt Laboratory, University of Oxford, UK). Lentiviral particles were generated by co-transfecting HEK293T cells with transfer, packaging, and envelope plasmids following established protocols (Tiscornia et al., 2006). Viral supernatants were harvested 48 hours post-transfection, filtered, and concentrated via ultracentrifugation to produce high-titre stocks.

### RNA isolation and qRT-PCR

Total RNA was extracted using the RNeasy Plus Mini Kit (Qiagen) according to the manufacturer’s instructions. Genomic DNA contamination was removed using the gDNA Wipeout buffer provided in the QuantiTect Reverse Transcription Kit (Qiagen), followed by cDNA synthesis from 1μg of total RNA. Quantitative real-time PCR (qPCR) was performed on a LightCycler® 2.0 Instrument using FastStart SYBR Green Master. Target gene expression was normalized to the housekeeping gene *hypoxanthine-guanine phosphoribosyltransferase* (*HPRT*). Relative mRNA fold changes were calculated using the 2^−ΔΔ*CT*^ method. Primer sequences are listed in the Supplementary Table S2.

### Statistical Analysis

Statistical analyses were performed using GraphPad Prism. Data distribution was assessed to determine the appropriate testing framework. For comparisons involving more than two groups, statistical significance was determined using a one-way ANOVA followed by Dunnett’s multiple comparisons test. For experiments involving two independent variables, a two-way ANOVA followed by Sidak’s multiple comparisons test was employed. Non-normally distributed data were analyzed using the Kruskal-Wallis test with Dunn’s multiple comparisons test. Direct comparisons between two independent groups were performed using an unpaired, two-tailed Student’s t-test. Data are presented as mean ± standard deviation (SD), and *p* < 0.05 was considered statistically significant.

## Disclosure and competing interests statement

Andrea Ballabio is a co-founder and shareholder of Casma Therapeutics. Frances M. Platt is a co-founder and shareholder of IntraBio. The remaining authors declare no competing interests.

## Funding

AB: the Italian Telethon Foundation, the Associazione Italiana per la Ricerca sul Cancro (IG-29203 to AB), the European Research Council (INCANTAR 101097752 to AB), the NIH (R01CA260205); EE: UCL-IoO studentship (JS) and Medical Research Council (grant number MR/V013882/1).

---

**Figure S1.**
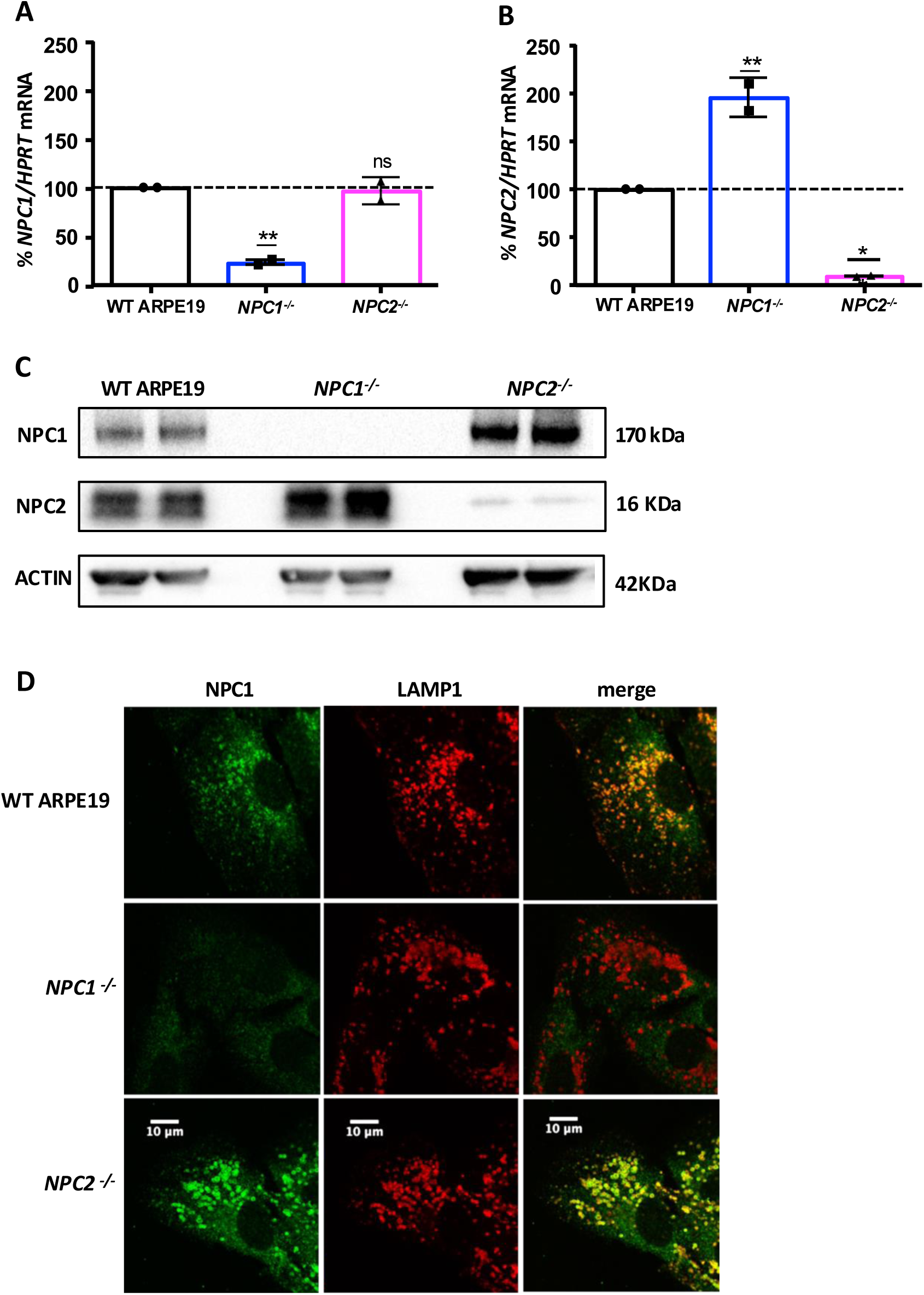
Validation of *NPC1* and *NPC2* knockout in *NPC1*⁻/⁻ and *NPC2*⁻/⁻ ARPE-19 cells. (A, B) qPCR analysis of (A) *NPC1* and (B) *NPC2* mRNA levels in WT, *NPC1*⁻/⁻, and *NPC2*⁻/⁻ ARPE-19 cells. Expression levels were normalized to *HPRT* and are presented as a percentage of WT levels. Primer sequences are provided in Table S2. (C) Western blot analysis of NPC1 and NPC2 protein levels in WT, *NPC1*⁻/⁻, and *NPC2*⁻/⁻ ARPE-19 cells. ACTIN served as a loading control. (D) Representative confocal microscopy images of WT, *NPC1*⁻/⁻, and *NPC2*⁻/⁻ ARPE-19 cells immunostained for LAMP1 (red) and NPC1 (green). Scale bar: 10 µm. Statistical analysis was performed using one-way ANOVA followed by Dunnett’s multiple comparisons test. Each sample was compared to the WT control. Data are presented as mean ± SD; ns = not significant; ** *p*<0.01.

**Figure S2.**
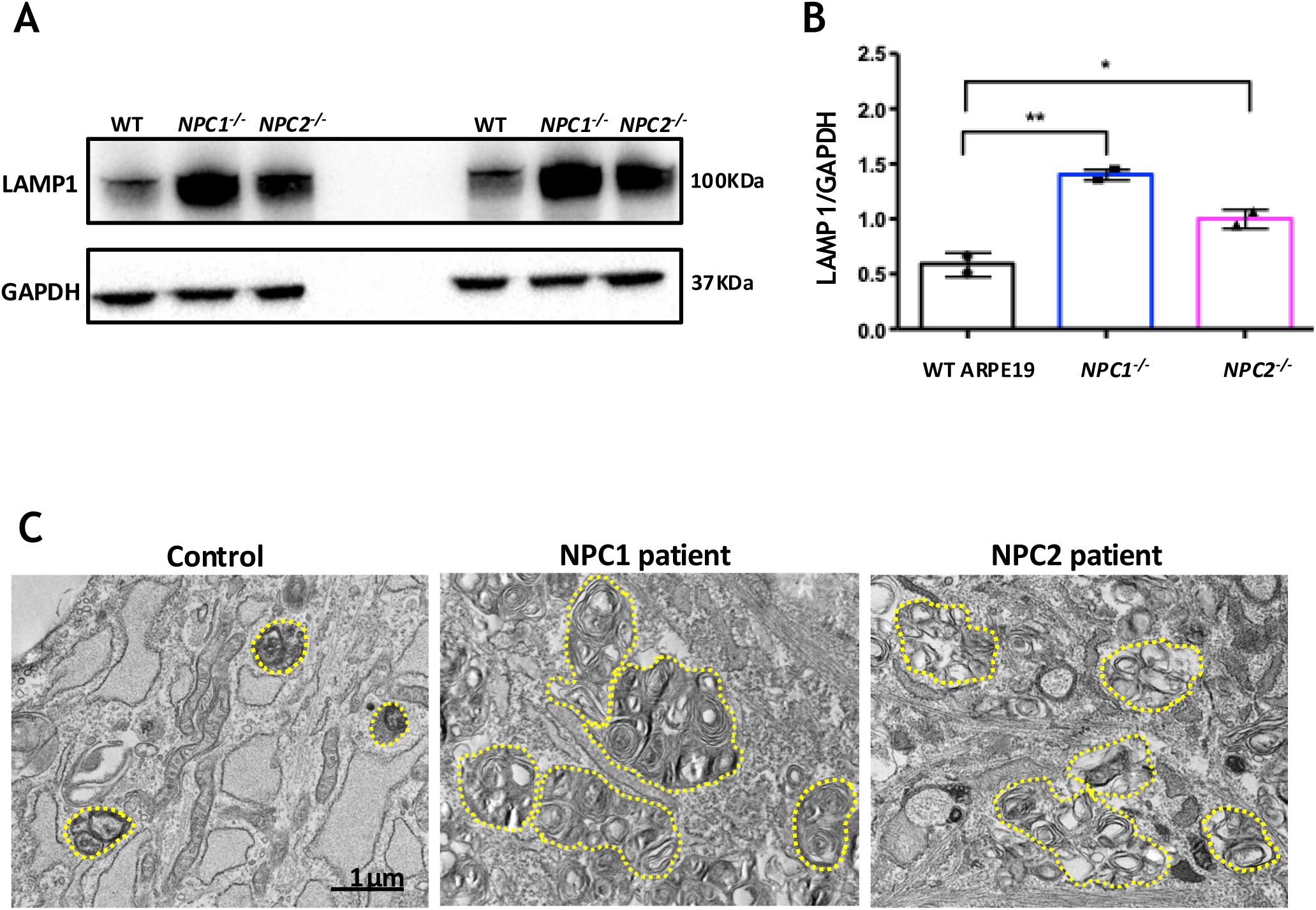
Increased LAMP1 expression in *NPC1*⁻/⁻ and *NPC2*⁻/⁻ ARPE-19 cells and lysosomal expansion in NPC patient fibroblasts. (A) Western blot analysis of LAMP1 expression in WT, *NPC1*⁻/⁻, and *NPC2*⁻/⁻ ARPE-19 cells. Total cell lysates were immunoblotted with antibodies against LAMP1 and GAPDH (loading control). (B) Densitometric quantification of the relative LAMP1 protein levels normalized to GAPDH (based on the representative blots shown in A). Statistical analysis was performed using one-way ANOVA followed by Dunnett’s multiple comparisons test. Each sample was compared to the WT control. Data are presented as mean ± SD; ** *p*<0.01, * *p*<0.05. Data are representative of 3 independent experiments. (C) Representative electron micrographs of skin fibroblasts derived from healthy donors (Control) or patients harbouring *NPC1* or *NPC2* mutations. Samples were processed for transmission electron microscopy (TEM) to assess organelle morphology. Individual lysosomes/late endosomes are delineated by dashed yellow outlines. Scale bar: 1μm. Morphometric analysis of endolysosomal parameters was performed using ImageJ (Fig 2D).

**Figure S3.**
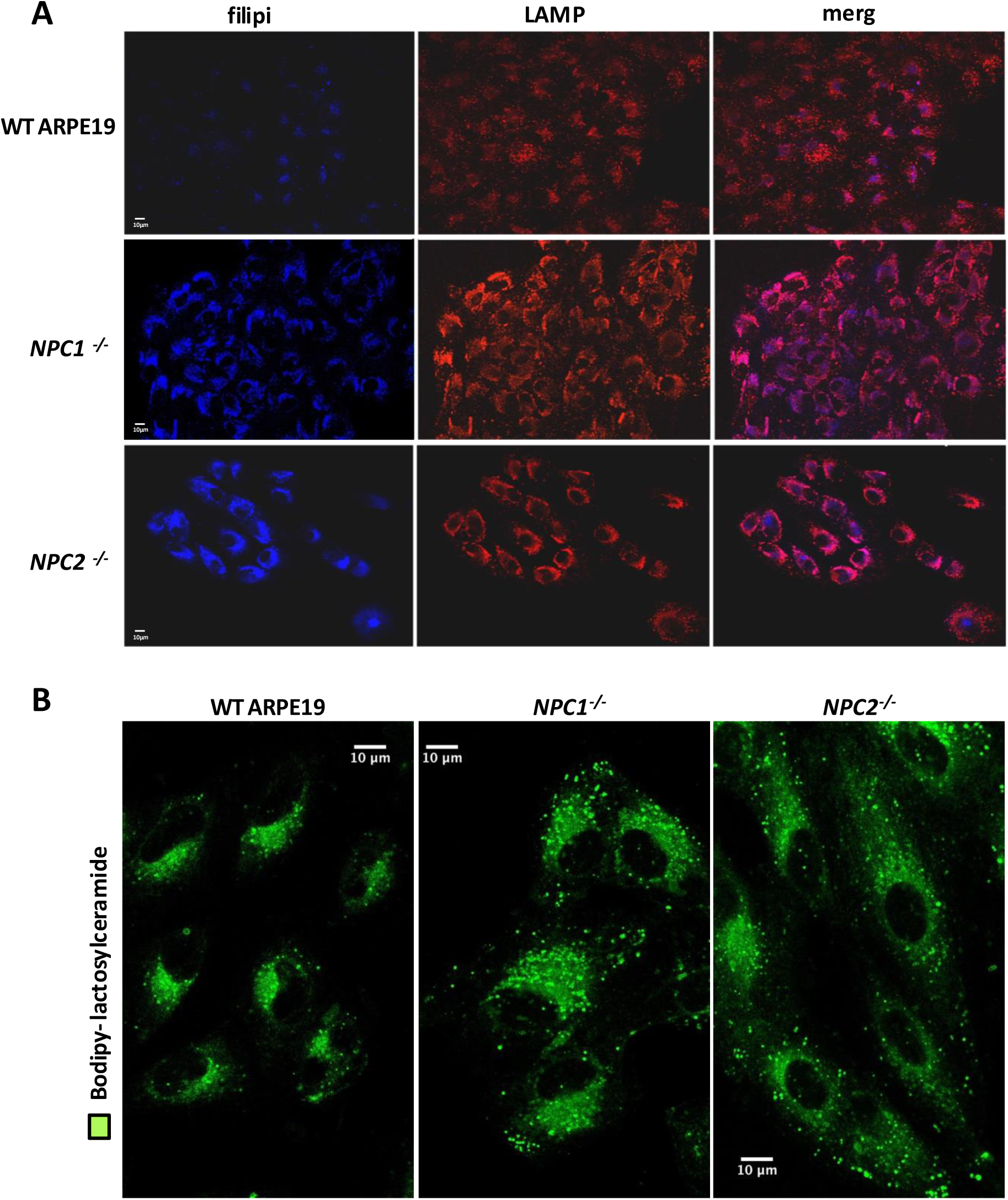
Characterization of lysosomal distribution and lactosylceramide trafficking. (A)Lysosomal distribution in WT, *NPC1*⁻/⁻, and *NPC2*⁻/⁻ ARPE-19 cells. Representative confocal microscopy images of WT, *NPC1*⁻/⁻, and *NPC2*⁻/⁻ ARPE-19 cells co-stained with filipin (blue) and LAMP1 (red). Scale bar: 10μm. In *NPC1*⁻/⁻ and *NPC2*⁻/⁻ cells, LAMP1-positive compartments exhibit a concentrated perinuclear distribution, whereas WT ARPE-19 cells show lysosomes dispersed throughout the cytoplasm. (B) Trafficking of BODIPY-LacCer in WT, *NPC1*⁻/⁻, and *NPC2*⁻/⁻ ARPE-19 cells. Live ARPE-19 cells were pulsed with 10μM BODIPY-LacCer (green), for 15 min at room temperature, followed by a 45-min chase at 37 °C to allow for endocytic trafficking. Cells were subsequently imaged by fluorescence microscopy. Images shown are representative of at least three independent experiments. Qualitative image assessment was performed by an observer blinded to the genotype.

